# Optimizing accuracy and depth of protein quantification in experiments using isobaric carriers

**DOI:** 10.1101/2020.08.24.264994

**Authors:** Harrison Specht, Nikolai Slavov

**Author notes:** Data & analysis code: scope2.slavovlab.net.

## Abstract

The isobaric carrier approach, which combines small isobarically-labeled samples with a larger isobarically-labeled carrier sample, is finding diverse applications in ultrasensitive mass-spectrometry analysis of very small samples, such as single cells. To enhance the growing use of isobaric carriers, we characterized the trade-offs of using isobaric carriers in controlled experiments with complex human proteomes. The data indicate that isobaric carriers directly enhances peptide sequence identification without simultaneously increasing the number of protein copies sampled from small samples. The results also indicate strategies for optimizing the amount of isobaric carrier and analytical parameters, such as ion accumulation time, for different priorities such as improved quantification or increased number of identified proteins. Balancing these trade-offs enables adapting isobaric carrier experiments to different applications, such as quantifying proteins from limited biopsies or organoids, building single-cell atlases, or modeling protein networks in single cells. In all cases, the reliability of protein quantification should be estimated and incorporated in all subsequent analysis. We expect that these guidelines will aid in explicit incorporation of the characterized trade-offs in experimental designs and transparent error propagation in data analysis.

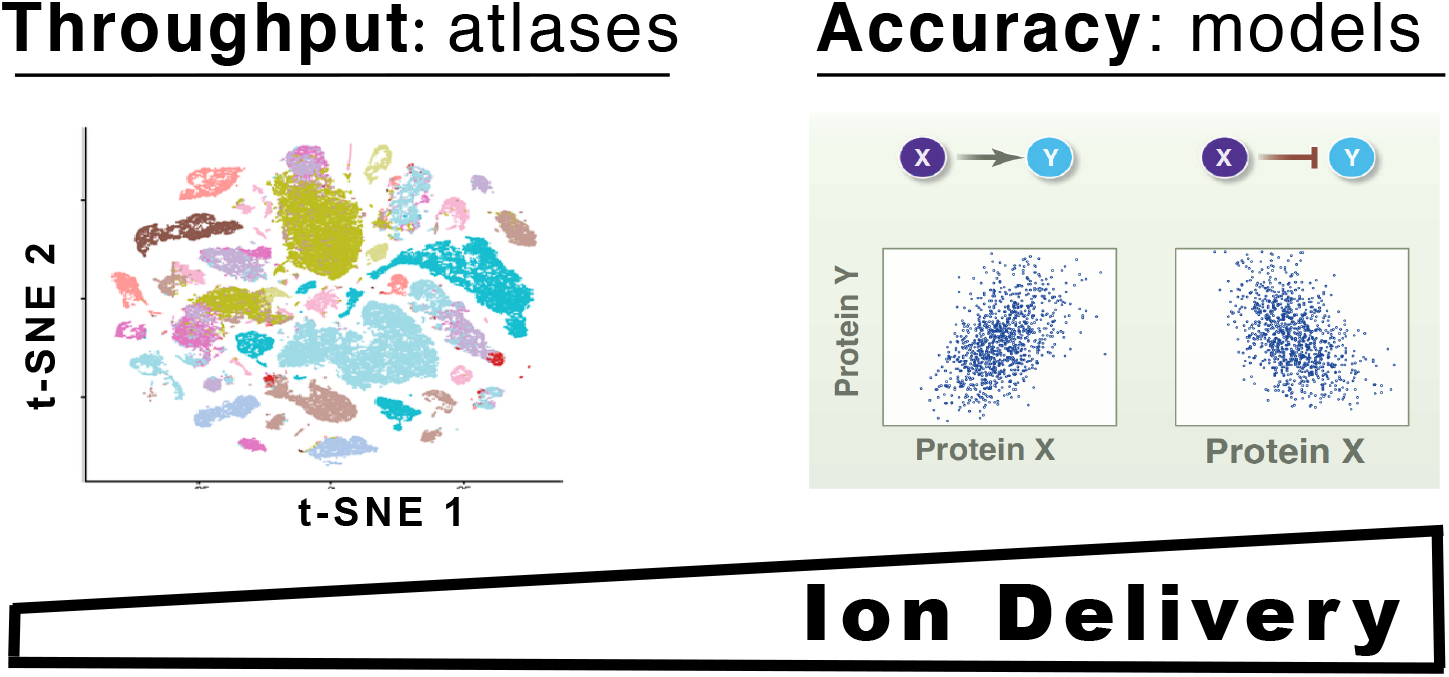

## Introduction

Today, mass spectrometry (MS) has become the most powerful method for analyzing proteins in bulk samples comprised of many cells.^1,2^ However, MS analysis of smaller samples, such as single cells, is more challenging because the ions analyzed by the MS detectors may be insufficient for accurate quantification and sequence identification.^3–8^ To mitigate these challenges, we introduced the *isobaric carrier* concept as part of Single Cell ProtEomics by Mass Spectrometry (SCoPE-MS),^9,10^ and the concept has been used by multiple laboratories as recently reviewed.^11^ The isobaric carrier approach employs tandem mass tags to label small samples of interest (e.g., proteomes of single cells) and a carrier sample (e.g., the proteome of 100 cells), and then combines all labeled samples to be analyzed together by tandem mass-spectrometry, as illustrated in Fig. 1.

**Figure 1.**
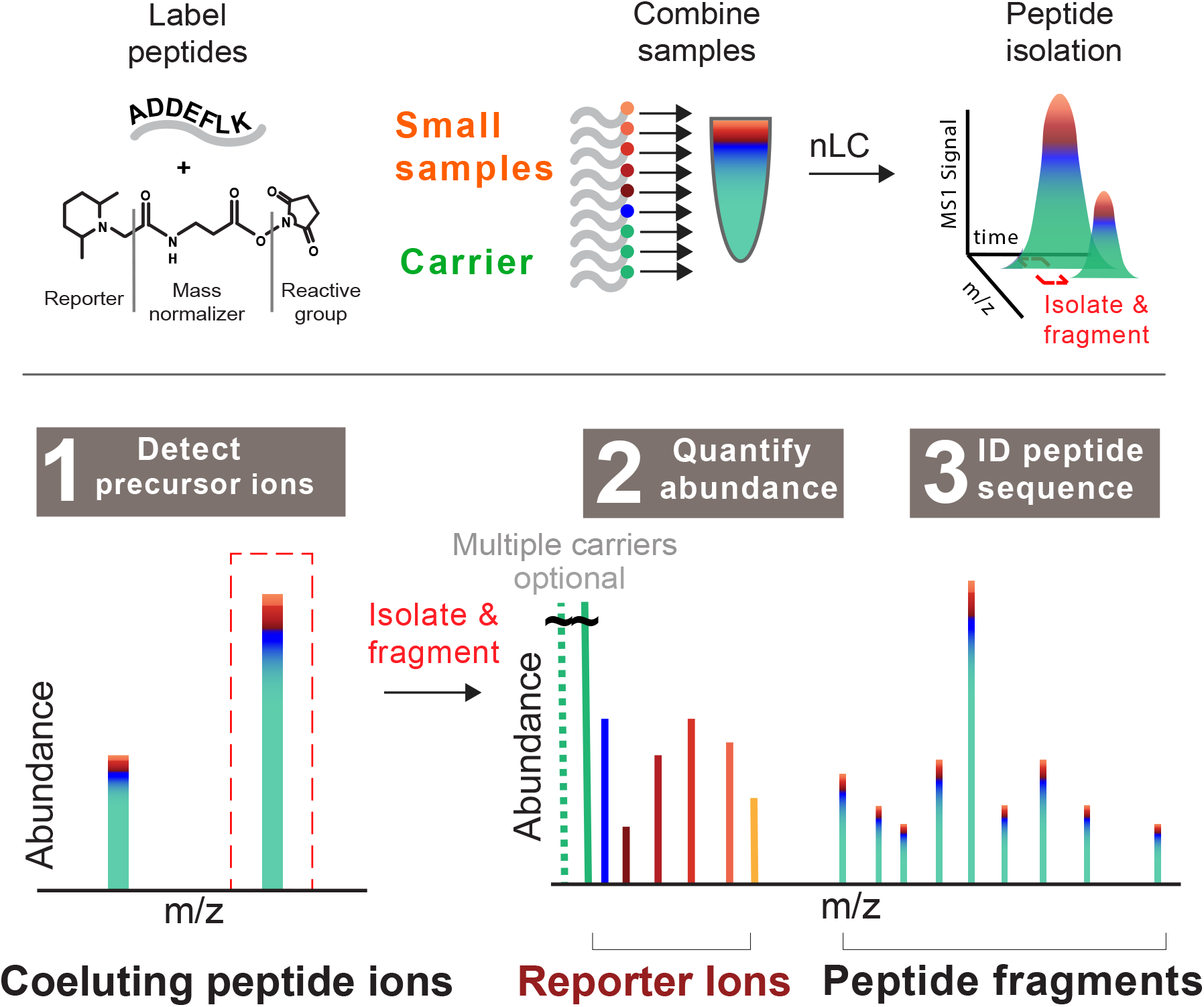
Schematic diagram of the isobaric carrier concept. The peptides from small samples and a from a larger (carrier) sample are labeled with isobaric tags, mixed and analyzed by tandem MS. Some sets, as the standards described in Table 1 may have more than 1 isobaric carrier samples. This approach increases the intensity of precursor ions (1), provides reporter ions for quantification (2) and facilitates sequence identifications based on the peptide fragments pooled across across all samples (3).

The isobaric carrier concept has been applied to single-cell protein analysis, detection of mutant proteoforms, phosphorylation and protein synthesis.^11^ This growing use of the concept motivated us to benchmark its benefits and limitations in controlled experiments and to extend the previously suggested approaches for optimizing ultrasensitive MS experiments.^12^ We explored the role of isobaric carrier in (i) facilitating the detection of precursor ions in MS1 survey scans and (ii) facilitating sequence identification by providing peptide fragment ions to MS2 spectra. These benefits must be balanced with possible adverse effects on quantification. Specifically, large levels of isobaric carriers may enable identifying peptides whose single-cell copies are insufficiently sampled by the MS detector to support accurate quantification.^11^

A fundamental and general challenge to single-cell analysis is sampling sufficient copies from each molecule type to support its reliable quantification. This remains, for example, a significant bottleneck for advanced single-cell RNA sequencing approaches.^13,14^ Nonetheless, some applications such as building cell-type and cell-state atlases can benefit from analyzing a large number of genes and single cells even if only a few copies are sampled from most messenger RNAs.^15^ Other applications, such as building biophysical models, demand accurate quantification and require sampling a larger number of copies per gene.^4,6,16^ Mass-spectrometry approaches for single-cell analysis have already demonstrated the ability to sample 20-fold more copies per gene compared to established single-cell RNA-seq methods.^17^

Sampling many copies per gene is challenging and often comes at the cost of decreased throughput both for single-cell RNA-seq^14,15^ and MS analysis.^4,17^ Thus, experimental designs should take into account the costs and benefits of each analytical strategy and choose whether to emphasize throughput or quantitative accuracy. This optimization has already been discussed^4,12,17^ but the trade-offs of varying the number of cells in the isobaric carriers remain incompletely characterized. Here we sought to empirically characterize these trade-offs and use the data to provide recommendations for experimental designs.

## Results

### Peptide sequence identification is a major bottleneck

To understand the major benefits of using the isobaric carrier, we began our analysis by considering the major challenges and bottlenecks for ultrasensitive MS analysis without using the isobaric carrier. Specifically, we aimed to evaluate the effects of the isobaric carrier on three stages of MS/MS data acquisition and analysis shown in Fig. 1: (1) detecting peptide-like features (precursor ions) during MS1 survey scans, (2) quantifying peptides from the small samples based on the reporter ions and (3) identifying peptide sequences from the fragment ions.

We evaluated the efficiency of each of these three steps for 1-cell and 100-cell samples analyzed by Cong *et al*..^18^ The data shown in Fig. 2a demonstrate that even for single cells, the mass spectrometer detected tens of thousands of peptide-like features and conducted MS2 scans on over 10,000 of these features. These numbers are comparable to the corresponding numbers for 100-cell samples and indicate that detecting precursor ions is not a limiting factor; indeed, the number of detected ions exceeds the number of MS2 scans that can be performed with the method setting.

**Figure 2.**
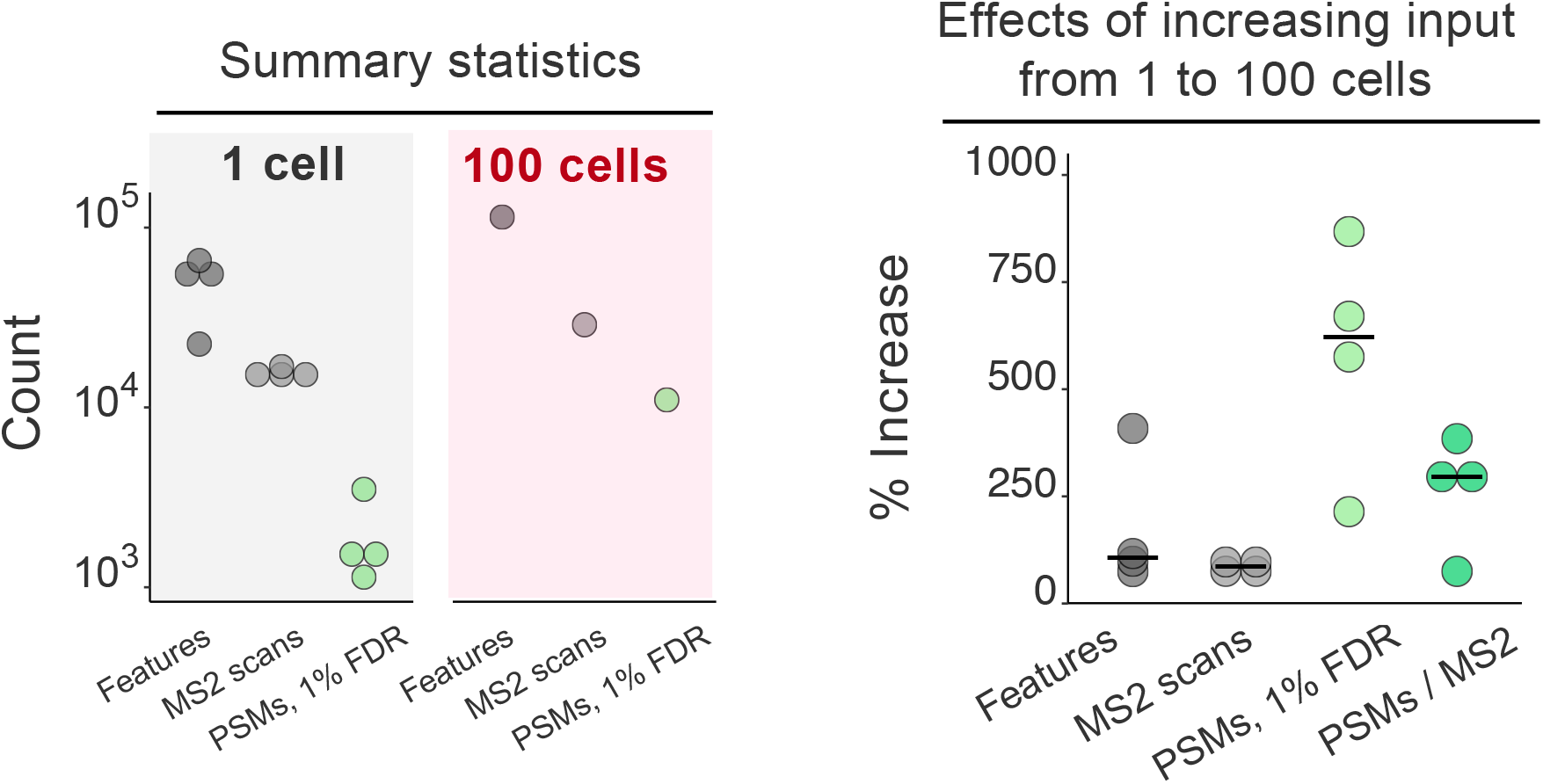
Increasing input from 1 to 100 cells primarily benefits the identification rate of MS2 spectra. Replicates of 1-cell and 100-cell HeLa samples were analyzed by label-free proteomics by Cong et al..^18^ (**a**) The number of peptide-like features (unique isotopic envelopes resolved with respect to m/z and retention time with charge ≥ +2), the number of MS2 spectra acquired, and the number of peptide-spectral-matches (PSMs) determined by MaxQuant at 1% false-discovery rate (FDR). (**b**) The percent increase in each metric from 1 to 100 cell input. The percent increase in PSMs and identification rate (PSMs / MS2: the number of PSMs at 1% FDR divided by the number of MS2 scans acquired) is greater than the percent increase in peptide-like features or MS2 scans acquired.

Thus, while the isobaric carrier can enhance the detection of precursor ions (stage 1 in Fig. 1), such enhancement is unlikely to significantly increase the number of MS2 scans conducted, because feature detection is not a limiting factor, as shown in Fig. 2; what is limiting is the speed of the mass spectrometer and the time for performing MS2 scans of the large number of identified peptide-like features. If needed, the feature detection can be further enhanced by increasing the accumulation times for survey scans, as can be afforded by narrower isolation windows, e.g., as implemented by BoxCar data acquisition.^19^

Despite the large number of MS2 scans, the rate of assigning confident peptide sequences is relatively low for the 1-cell samples and increases by over 250 % for the 100-cell sample, Fig. 2b. This increase is likely due to increased diversity and abundance of observed peptide fragment ions for the 100-cell sample. Thus, obtaining enough peptide fragments for confident sequence identification is a major bottleneck in the analysis of small samples, such as individual cells. The peptide fragments provided by the isobaric carrier (as illustrated in Fig. 1) can help overcome this bottleneck.

### Theoretical expectations

While the isobaric carrier can bolster peptide sequence identification, it does not increase the number of ion copies sampled from the small-sample peptides. Therefore, the isobaric carrier may support confident peptide identification even when the peptide copies sampled from the small samples are insufficient to support reliable quantification.

Before empirically examining the effects of increased amounts of isobaric carriers, we consider the theoretically expected effects as sketched in Fig. 3. Without an isobaric carrier, the rate of accumulating ions from the small samples is low and thus the ion accumulation is likely to use the maximum allowed accumulation time before reaching the operator-defined automatic gain control (AGC) target. As the amount of the isobaric carrier increases, the rate of ion accumulation increases as well, and accumulated ions begin to reach the AGC target before the maximum fill time. Thus, the fill time begins to decrease resulting in decreased sampling of ions from the small samples. The shorter fill times can decrease the cycle times and thus increase the number of analyzed peptides, which can be sent for MS2 scans and reliably identified thanks to the fragment ions originating from the isobaric carrier and the small samples.

**Figure 3.**
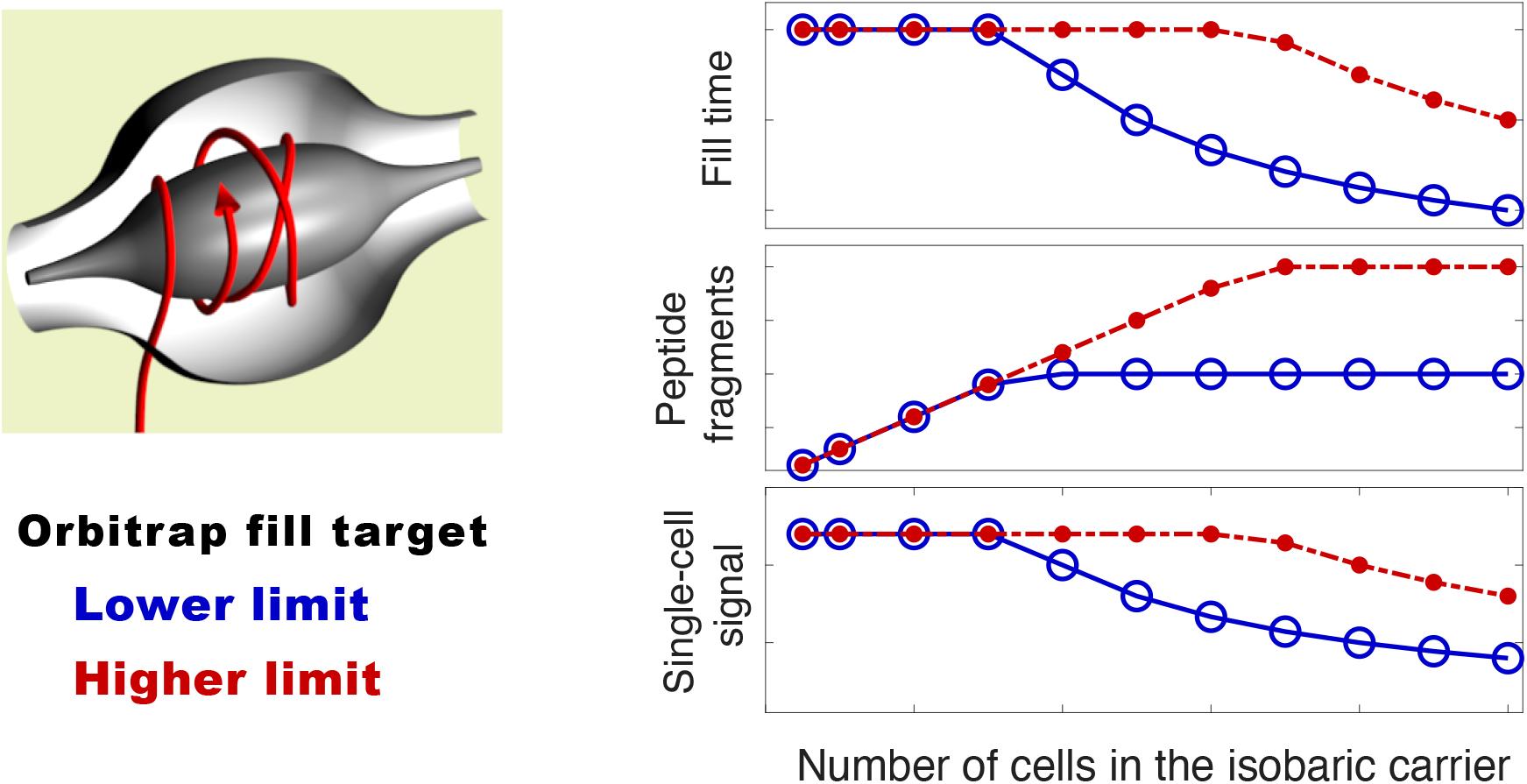
Theoretically-expected effects of increased isobaric carrier. Increasing the number of cells in the isobaric carrier increases the rate of accumulating peptide fragments for MS2 analysis. When the target AGC is reached, accumulation of ions stops, which may increase the speed of the analysis at the expense of decreased sampling of peptides from the small samples.

Increasing the AGC target increases the size of the isobaric carrier at which accumulation times (and ion sampling from the small samples) begin to decrease, Fig. 3. Thus, higher AGC targets may increase the copy number of ions sampled from the small samples at the expense of more time needed for MS2 analysis of each peptides. This theoretical example illustrates a clear trade-off that next we explore empirically in controlled experiments.

### Effects of isobaric carriers on peptide identification

To empirically benchmark the effects of the isobaric carrier, we created bulk standards that, when diluted 100-fold, model SCoPE2 standards with two carrier channels and 6 single-cell channels, as shown in Table 1. The sets have isobaric carriers with sizes ranging from 100-cell to 800-cell equivalents, Table 1. Standards with larger carriers result in identifying more peptides (Fig. 4), including additional less abundant peptides not identified in the standards with smaller carrier. The larger carriers might support much higher peptide identification rates with different parameters (faster ion accumulation and MS2 scans), but we kept these parameters constant across all experiments to allow for controlled comparisons. Thus, the 256ms MS2 transient times required for 70k resolution do not allow taking advantage of the short accumulation times for large-size carriers and low AGC target, about 40ms for the 800-cell carrier as shown in Fig. 5a; see DO-MS reports. To demonstrate the potential for identifying more peptides, we reduced the MS2 transient times (by reducing the resolution to 35,000), which increased the number of MS2 scans and peptide identifications at the low AGC target, as shown by the dotted curve in Fig. 4a.

**Table 1.** Standards with variable amounts of isobaric carriers. We prepared a series of diluted standards approximating SCoPE2 sets with different amounts of cells in their isobaric carriers. U stands for U937 (a cell line of monocytes) and J stands for Jurkat cells (a cell line of T-cells). The number in front of the letter indicates the number of cell equivalents in each sample injected for MS analysis.

| Carrier size | 126 | 127N | 127C | 128N | 128C | 129N | 129C | 130N | 130C | 131N | 131C | 132N | 132C | 133N | 133C | 134N |
| --- | --- | --- | --- | --- | --- | --- | --- | --- | --- | --- | --- | --- | --- | --- | --- | --- |
| 100-cell | 50J | 50U | - | - | 1J | 1U | 1J | 1U | 1J | 1U | 1J | 1U | - | - | - | - |
| 200-cell | 100J | 100U | - | - | 1J | 1U | 1J | 1U | 1J | 1U | 1J | 1U | - | - | - | - |
| 300-cell | 150J | 150U | - | - | 1J | 1U | 1J | 1U | 1J | 1U | 1J | 1U | - | - | - | - |
| 400-cell | 200J | 200U | - | - | 1J | 1U | 1J | 1U | 1J | 1U | 1J | 1U | - | - | - | - |
| 600-cell | 300J | 300U | - | - | 1J | 1U | 1J | 1U | 1J | 1U | 1J | 1U | - | - | - | - |
| 800-cell | 400J | 400U | - | - | 1J | 1U | 1J | 1U | 1J | 1U | 1J | 1U | - | - | - | - |

**Figure 4.**
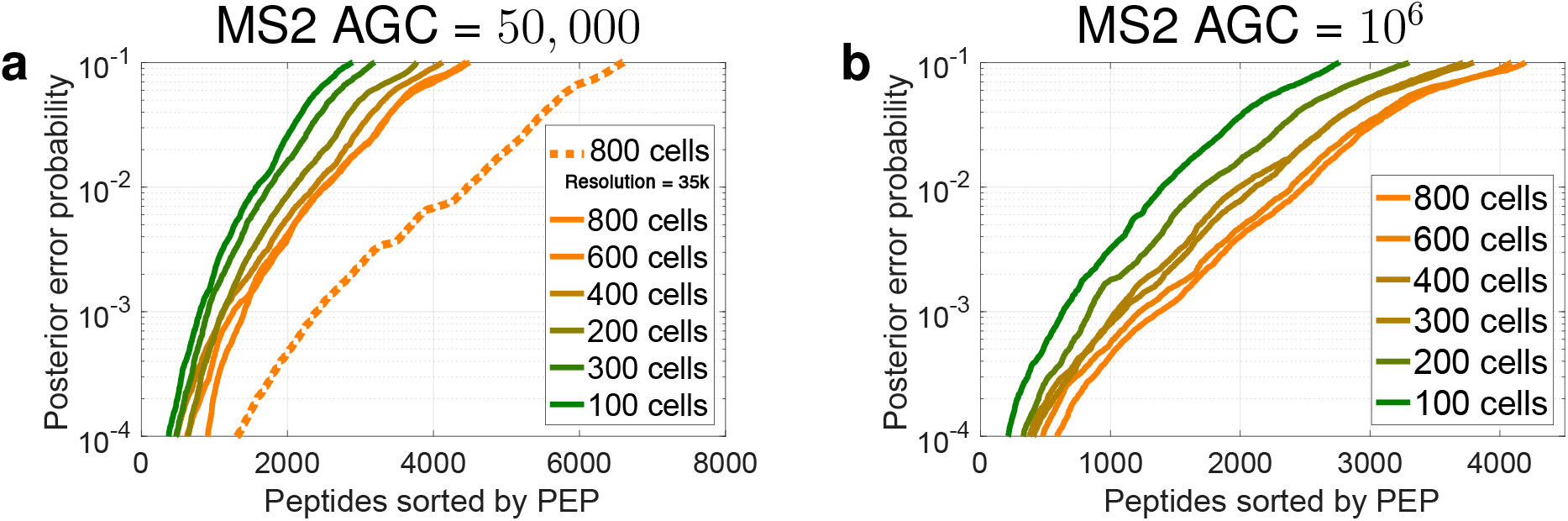
Number of identified peptides. Increasing the number of cells in the isobaric carrier results in larger number of confidently identified peptides at any level of confidence as quantified by the posterior error probability (PEP). This trend is observed both with the low MS2 AGC target (50,000) shown in (**a**) and with the high MS2 AGC target (1,000,000) shown in (**b**). The MS2 scans of the run displayed with a dotted curve were performed at 35k resolution to demonstrate the potential advantage of low AGC for scanning and identifying more peptides. The MS2 scans of all other runs were performed at 70k resolution. The PEPs are estimated by MaxQuant using only the mass spectra^20^ and do not include additional features, such as retention time.^3^ These plots are standard component of the DO-MS reports,^12^ and the full reports are included in the supplemental data and at scope2.slavovlab.net.

**Figure 5.**
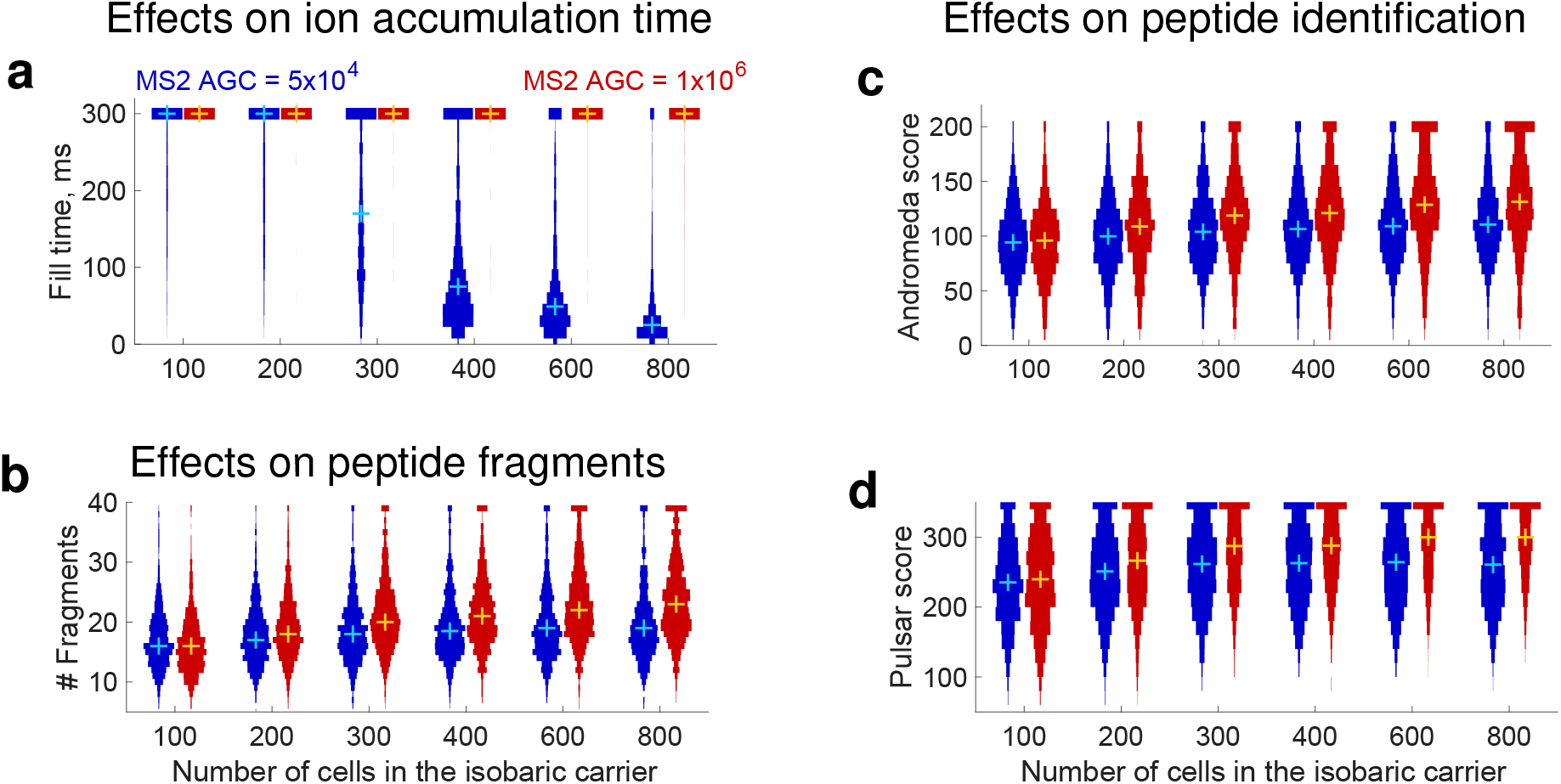
Effects of increasing the size of the isobaric carrier on peptide accumulation and sequence identification. (**a**) Distributions of MS2 accumulation times for all peptides identified across all displayed experiments with standards described in Table 1. (**b**) Number of peptide fragments detected per PSM for all peptides that are identified in each experiment. For visualization purposes, peptides with more than 39 fragments were set to have 39 fragments. The confidence of sequence identification for peptides identified across all experiments is displayed as distributions of scores computed either by Andromeda (**c**) or by Pulsar (**d**). These scores quantify the confidence of peptide-spectrum-matches.^22,23^ Andromeda scores exceeding 200 were set to 200 for visualization purposes. The distributions shown here can be generated by DO-MS (DO-MS; Data-driven Optimization of MS is software freely available at do-ms.slavovlab.net) and can be used to evaluate the regime of analysis for any particular set of experiments.^12^ To enable controlled comparisons, the distributions show data only for the subset of peptides identified across all levels of isobaric carriers.^12,21^ The plus marks denote medians.

If this compositional difference is not taken into account, the distributions of peptide quantities cannot be meaningfully compared; comparisons between distributions containing different subsets of peptides may lead to substantial biases. This phenomenon is well know as *missing not at random* or *nonignorable missingness*^21^ and can case erroneous interpretations of proteomics data.^12^ To avoid such problems, we controlled for different peptide compositions by focusing the analysis only on the subset of peptides identified across all samples.

As theoretically expected for Fig. 3, increasing the carrier amount results in reaching the low AGC target before the maximum fill times, Fig. 5a: for carrier channels exceeding 300 cells, we observe that the low AGC target may be reached before the max fill time. However, the high AGC target is not reached by most peptides within 300 ms even with an 800-cell isobaric carrier, Fig. 5a.

As the amount of isobaric carrier increases, some peptides begin to reach the low AGC target (50,000), at about the 300-cell carrier. For larger carriers, the accumulated peptide fragments and the confidence of peptide identification remain constant Fig. 5b,c,d, while the small-sample signal decreases proportionately to the decreased fill times Fig. 6a. High AGC target results in different trends: the accumulated peptide fragments and the confidence of peptide identification increase with the carrier size Fig. 5b,c,d, while the small-sample signal remains constant Fig. 6a. This effect of an increased MS2 AGC target is consistent with previous observations,^24^ and previously used analytical parameters for SCoPE-MS and SCoPE2 analysis have corresponded to the high AGC target regime.^10,17^ In this regime, the isobaric carrier does not limit MS2 accumulation times and ions are accumulated for the maximum time allowed as shown in Fig. 5a.

**Figure 6.**
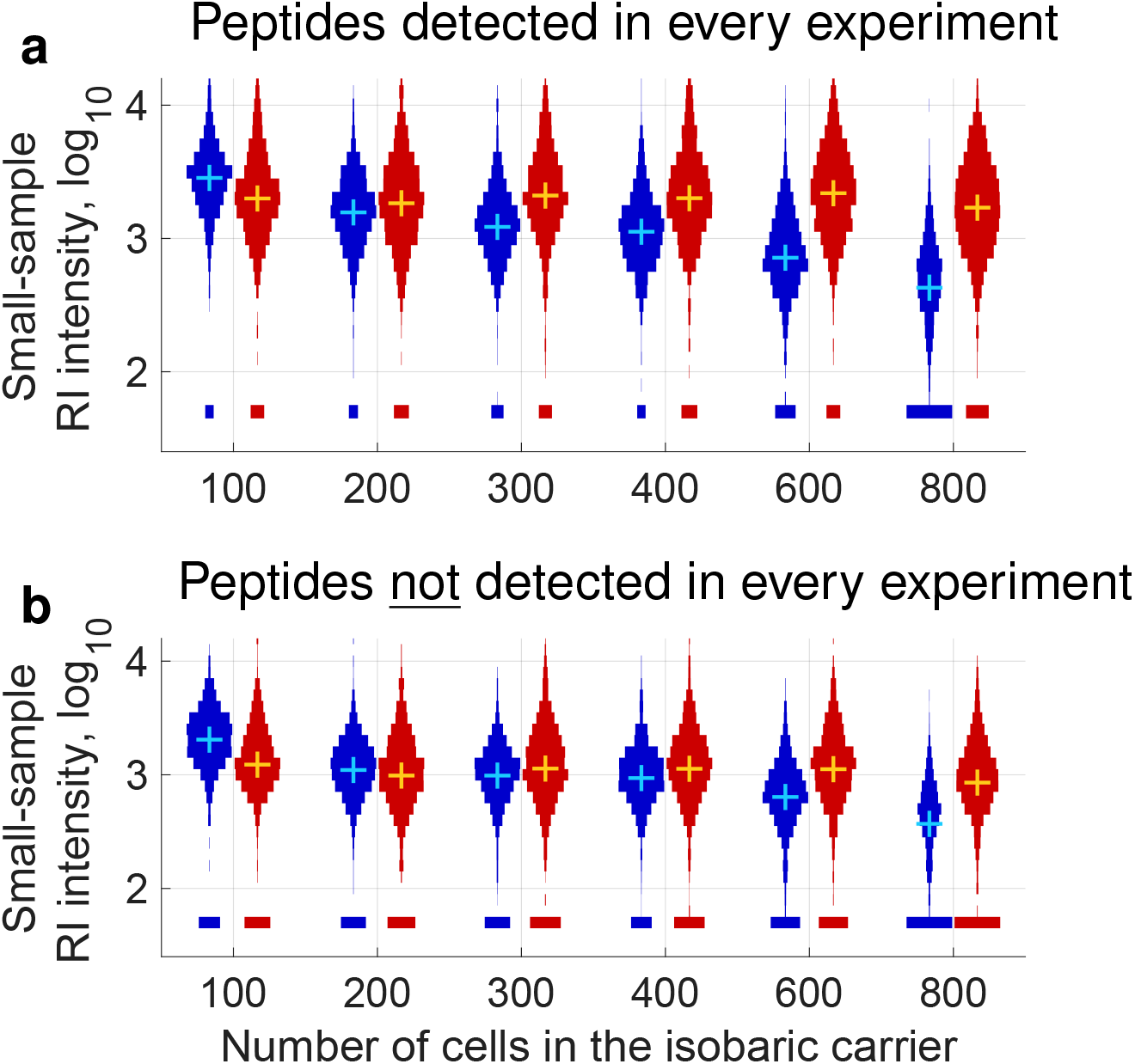
Effects of increasing the size of the isobaric carrier on the reporter ion intensities in the small samples. Both panels shows distributions of reporter-ion (RI) intensities from the small samples of the standards described in Table 1. As in Fig. 5, the blue distributions correspond to MS2 AGC = 50,000, and the red distributions correspond to MS2 AGC = 1,000,000 (**a**) Only the RI intensities for peptides identified across all experiments are shown to allow for a well-controlled comparison.^12,21^ (**b**) Only the RI intensities for peptides not identified across all experiments are shown to evaluate whether some of these peptides have high enough RI intensity to be quantifiable. The means and medians of these distributions cannot be meaningfully compared because of nonignorable missing data.^21^

In addition to comparing the distributions of small-sample RI intensities for the peptides detected across all carrier levels (Fig. 6a), we also compared the small-sample RI intensities for the peptides detected only in some samples, Fig. 6b. While this comparison is not well-controlled, it allows to evaluate whether the additional peptides detected with larger isobaric carriers are sampled with sufficient copy numbers to allow quantification in the small samples. These additional peptides are likely less abundant and sampled with fewer copies. Indeed, the distributions of small-sample RI intensities tend to decrease with the increase of isobaric carrier for both MS2 AGC targets, Fig. 6b. However, many of these additional peptides have high enough RI intensities to have the potential to support quantification, especially with the higher AGC MS2 target, Fig. 6b.

### Effects of isobaric carriers on peptide quantification

The number of ion copies sampled per peptide is an important determinant of quantification accuracy.^4,17^ Thus, the decreased ion copy number sampling at low MS2 ACG target and high carrier are likely to adversely affect quantification accuracy. However, the magnitude of this effect is unclear because other factors contribute significantly to the accuracy of quantification, such as the effi-ciency of sample preparation and the unintentional coisolation of multiple precursor ions for MS2 analysis. Indeed, the same ion sampling efficiency results in more accurate protein quantification in standards diluted to single-cell levels than in single-cell SCoPE2 sets.^17^ This fact underscores the importance of sample preparation recently reviewed by Kelly^7^ and demands the use of direct benchmarks for quantification accuracy.

To benchmark relative quantification in the single-cell channels, we used the quantification derived from the two carrier channels present in each set, Table 1: We correlated the peptide foldchanges estimated in the small samples to the corresponding peptide fold-changes estimated from the isobaric carrier channels, Fig. 7a. The results indicate that for the high AGC target (10^6^), the correlations remain relatively constant as the size of the carrier increases. In stark contrast, the correlations decline for the low AGC target, Fig. 7a, in parallel to the decline of reporter ions intensities shown in Fig. 6a. Likewise, we correlated the peptide fold-changes estimated in the small samples to each other in Fig. S1. The same trend is observed as when correlating to fold changes estimated from the isobaric carrier channel. These results indicate that sampling fewer copies from single cells is not simply a function of a large isobaric carrier and hardware limitations; rather, it is a reflection of analytical parameters that favor speed of analysis (e.g., lower AGC) and number of identified peptides over sampling many ion copies. While this regime results in less quantitative data, the increased throughput might offer worthwhile advantages for some applications, such as building single-cell atlases.^15^

**Figure 7.**
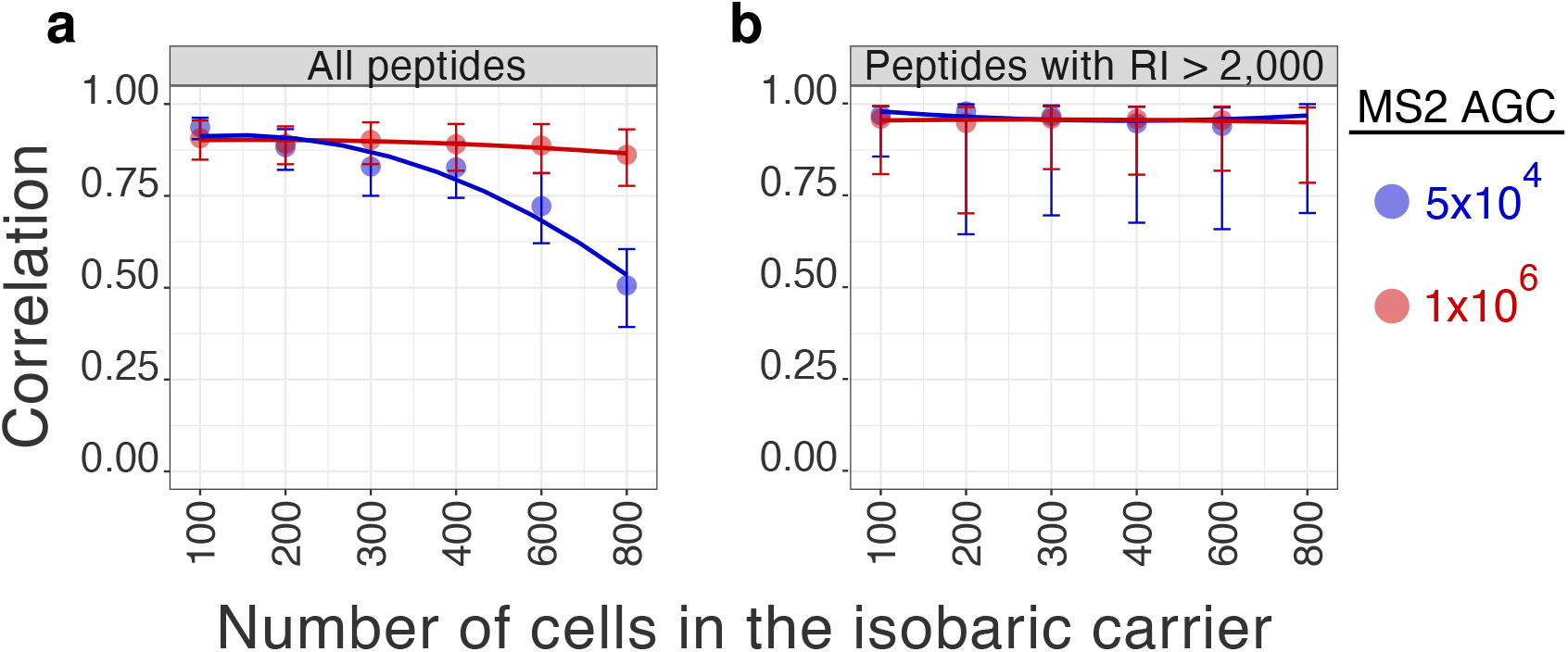
Effects of increasing the size of the isobaric carrier for two target levels of peptide quantification. (**a**) Correlations between the protein fold-changes (between monocytes and T-cells) estimated from the small samples and the carrier samples. All peptides quantified across all samples at 1% FDR are used for this analysis. For the lower AGC series, there were 966 unique peptides and 382 unique proteins quantified in every experiment. For the higher AGC series, there were 982 unique peptides and 349 unique proteins quantified in every experiment. (**b**) Correlations between fold changes as in panel (a) but only for peptides whose reporter ion intensities in the small samples are larger than 2,000. In both panels, error bars represent the 90% confidence intervals computed from resampling subsets of the data. For the lower AGC series, there were 725 unique peptides and 298 unique proteins quantified in every experiment. For the higher AGC series, there were unique 657 peptides and 240 unique proteins quantified in every experiment.

To further test the interpretation that quantitative accuracy is lower because of decreased sampling of ion copies, we evaluated the quantitative accuracy for the subset of peptides having reporter ion intensities above 2,000. The high correlations between fold change vectors (*ρ* > 0.9) indicate good accuracy for all carrier sizes and both AGC targets, Fig. 7b. These result affirm that in our experiments, quantitative accuracy depends on sampling enough ion copies from the small samples. Furthermore, the results emphasize that the sampling efficiency and quantitative accuracy is peptide and protein specific; a single experiment contains both well quantified and poorly quantified proteins. Therefore, we see great benefit in estimating the reliability of quantification for each protein and using this reliability for further quantitative analysis of the data, e.g., for correcting estimates of the fraction of explained variance.^25^

The data presented so far illustrates the effects of parameters that indirectly alter the MS2 accumulation time. To further demonstrate these effects, we directly limited MS2 accumulation times to 100, 200, 300, and 600 ms. The results demonstrate that longer accumulation times result in higher confidence of peptide identification (Fig. 8a), more samples ions from the small samples (Fig. 8b), improved quantification (Fig. 8c) and decreased missing data (Fig. 8d). Importantly, the trends in Fig. 8c-d demonstrate that the improvements in the quantification of the small samples (lysates diluted to single-cell level) saturate at about 300ms. This corresponds to the default accumulation time used in this and in previous SCoPE2 analysis.^17^ Different parameters (and less abundant proteins) may benefit from longer accumulation times, and we recommend using this analysis (available via DO-M^S12^) to establish optimal accumulation times.

**Figure 8.**
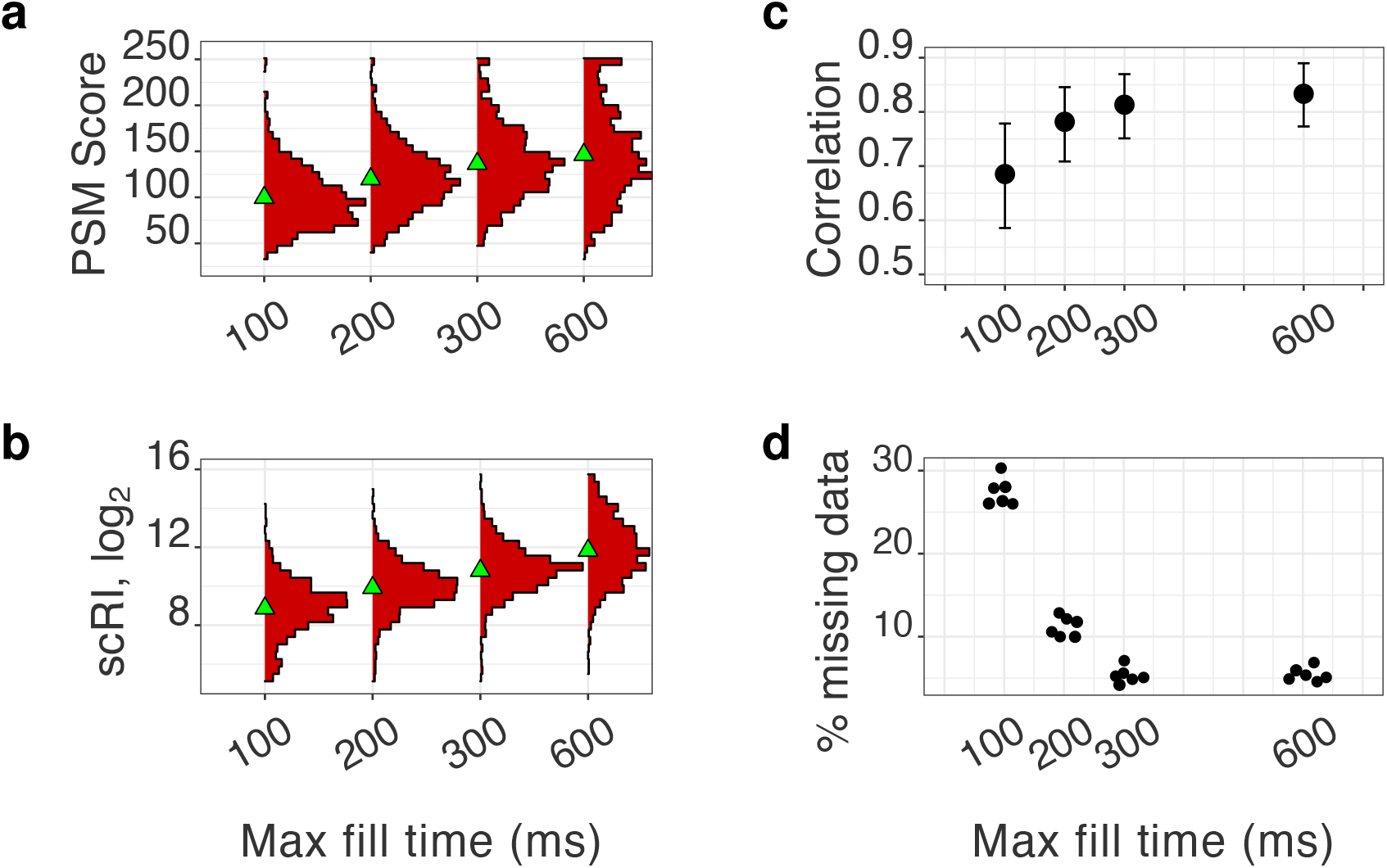
Effects of increasing the max MS2 fill time from 100ms to 600ms for a standard with a 100-cell isobaric carrier. All peptides quantified across all samples at 1% FDR are used for this analysis. (**a**) The confidence of sequence identification for peptides is displayed as distributions of scores computed by Andromeda.^22^ (**b**) Distributions of reporter-ion intensities from the small samples (scRI) of a standard with a 100-cell isobaric carrier across all max fill times. (**c**) Correlations between the protein fold-changes (between monocytes and HEK-293) estimated from the small samples and the isobaric carrier samples. (**d**) Fraction of missing reporter-ion intensities from the small samples as a function of max fill time.

## Discussion

A fundamental challenge to analyzing very small samples, such as individual mammalian cells, is sampling sufficient copies from each molecule to support its reliable quantification. While singlecell RNA-seq methods have generated much useful data without overcoming this challenge,^14,15^ some applications require accurate quantification that can be achieved only by sampling enough ion copies.

The data in Fig. 6 indicate that the sampling challenge is not created by the isobaric carrier approach but may be exacerbated by it in two ways. First, the isobaric carrier approach enables identifying the sequence of peptides that may be insufficiently sampled to be reliably quantified in the small samples. Second, poor experimental design (such as insufficient ion accumulation time or very large carrier amount and low limit on total ion accumulation) may reduce the copy number of sampled ions, and thus undermine quantification. The data presented here indicate that both pitfalls can be overcome by estimating the sampling efficiency for each peptide, and then using for further quantification peptides with enough sampled copies to support reliable quantification, Fig. 7b. We suggest that experimental designs optimize the isobaric-carrier amount and the ion accumulation times to reflect the relative priorities of throughput (number of proteins and cells analyzed per unit time) and quantification accuracy (copy number of ions sampled). The results should emphasize sampling efficiency and reliability estimates for each quantified protein rather than merely the number of identified peptides and proteins.

The data presented here illustrate a fundamental trade-off between throughput (number of cells and proteins analyzed) and the number of copies sampled per peptide from the small samples. The cornerstone of this trade-off is the fact that the isobaric carrier does not amplify (boost) the reporter ion intensities of peptides from the small samples (e.g., single cells). Therefore, delivering sufficient number of ion copies from small samples is essential for accurate quantification, as previously suggested^4,11,17^ and demonstrated here in Fig. 7. Thus, two modalities of ultrasensitive MS analysis emerge: analysis aiming to maximize quantitative accuracy must increase the delivery of analytes to the MS detector;^11^ analysis aiming to maximize throughput must reduce the time spent per analyte. To balance these competing priorities, we offer the following guidelines for LC-MS/MS experiments employing the isobaric carrier concept.

### Guidelines

1. **The size of the isobaric carrier should reflect the project priorities.** The data presented here reinforce previous suggestions that isobaric carriers that are about 200-fold larger than the small samples provide most of the needed increase in peptide fragments to enhance sequence identification without adverse affects on quantification.^9,17^ Larger carrier sizes can further benefit peptide sequence identification (Fig. 4 and Fig. 5) but will result in identifying additional less abundant peptides that are sampled with fewer copy numbers in the small samples (Fig. 6). If more than one carrier is used, optimization depends on the sum of all cells in all carriers.
2. **Evaluate whether the isobaric carrier affects peptide sampling.** We recommend estimating whether the carrier levels and AGC target result in reduced accumulation times and sampling of proteins from the small sample. This can be visualized by plotting the distributions of accumulation times and reporter ion intensities as illustrated in Fig. 5 and Fig. 6. These distributions can be automatically generated by DO-MS.^12^
3. **Estimate the reliability of quantification for each protein.** Estimate the sampling error on a per-protein basis as previously demonstrated,^17^ benchmark protein quantification as shown in Fig. 7, or estimate the reliability of quantification based on the consistency of the quantification of peptides originating from the same protein as demonstrated by Franks *et al*..^25^ If the reliability of quantification is limited by counting noise, improved sampling can increase reliability. However, if the reliability is limited by other factors, such as sample preparation, improved copy-number sampling may have limited benefits.
4. **Incorporate estimates of reliability in all subsequent analysis.** Data points should be weighted based on their reliability, with weights proportional to the reliability. Use error propagation methods to reflect the noise in the final results. For example, correlations between noisy variables can be divided by the corresponding reliability to estimate the fraction of explained variance independent from the noise.^25^

### Coisolation in the context of isobaric carrier

As with any approach using tandem mass-tags, quantification of samples employing isobaric carriers can be severely undermined by coisolating ions. Therefore, methods employing isobaric carriers should minimize coisolation by using narrow isolation windows to sample ions for MS2 scans, and by aiming to sample them when the abundance of the ions is highest (the apex of its elution peak), and by employing high performance chromatography that affords sharp elution peaks with minimal overlaps. These approaches have significantly decreased coisolation and improved quantification in SCoPE2 analysis when compared to SCoPE-MS.^17^

The degree of coisolation can vary across a set of isobarically labeled samples analyzed by the same LC-MS/MS run. This variation is due to the fact the coisolating analytes can vary in abundance across different samples. The degree of this variation depends on the samples analyzed together and is larger for samples that differ more, such as an isobaric carrier sample.

## Conclusion

Our data and analysis suggest that a principal benefit of the isobaric carrier is enhanced peptide sequence identification. Increasing the amount of isobaric carriers may allow faster peptide analysis and identification rate, but the associated decrease in accumulation times decreases the copy numbers of ions sampled from the small samples. Thus, the amounts of the isobaric carrier must reflect the balance of peptide sampling and depth of quantification that are best suited for the analysis. Then, the quantification errors should be estimated and reflected in any subsequent analysis.

## Materials and Methods

### Sample preparation

All analysis used samples corresponding to 100x standards prepared in bulk and diluted 100-fold to model single-cell SCoPE2 set. These standards were prepared as previously described.^17^ Specifically, U937 and Jurkat cells were collected from exponentially growing cultures, washed twice with cold PBS, and counted under hemocytometer to estimate cell density. The cells were then lysed in HPLC-grade water according to the mPOP protocol:^26^ a 15 min freeze step at −80 °C, followed by a 10 min heating step at 90 °C. Following lysis, samples were digested at 37 °C for 3 h with 10 ng/*μ*L of Promega Trypsin Gold in 100 mM TEAB. The bulk digested material was then serially diluted and labeled to generate the 16-plex design schemes shown in Table 1. These standards were prepared as 100x bulk samples with total volume of 100 *μ*L and 1% (1 *μ*L) of each bulk sample was then injected for MS analysis to simulate a single SCoPE2 experiment with two carrier channels of the indicated cell number and six single-cell channels, Table 1. No biological replicates were prepared of these bulk samples.

### MS analysis

The standards were analyzed with the MS methods used for SCoPE2 samples.^17^ Specifically, all samples were separated on an IonOpticks Odyssey (ODY-25075C18A) analytical columns with a 20 cm length and 75 *μ*m inner diameter. It was run by a Dionex RSLC3000 nano LC. All samples were analyzed by a Thermo Scientific Q-Exactive mass spectrometer. RAW files were searched using MaxQuant 1.6.7.0.^27^ The human SwissProt FASTA database (20,375 entries, downloaded 8/22/2020) was used for searching data from U-937 and Jurkat cells. Trypsin was specified as the digest enzyme, and a maximum of two missed cleavages were allowed for peptides between 5 and 26 amino acids long. Methionine oxidation (+15.99491 Da) and protein N-terminal acetylation (+42.01056 Da) were specified as variable modifications. TMTpro (+304.207145) was specified as fixed modification on lysine and peptide N-terminus. RAW files were also searched using SpectroMine 2.1.200828.47784.^23^ The human SwissProt FASTA database (20,375 entries, downloaded 8/22/2020) was used for searching data from U-937 and Jurkat cells. Trypsin was specified as the digest enzyme, and a maximum of two missed cleavages were allowed for peptides between 7 and 52 amino acids long. Methionine oxidation (+15.99491 Da) and protein N-terminal acetylation (+42.01056 Da) were specified as variable modifications. TMTpro (+304.207145) was specified as fixed modification on lysine and peptide N-terminus. MaxQuant results from Cong *et al*. were downloaded from the publication’s supplementary material.^18^

### Data availability

All data sets associated with this manuscript have been deposited at massIVE with ID: MSV000086004 and at scope2.slavovlab.net/mass-spec/Isobaric-carrier-optimization.

### Data analysis and visualization

Data analysis followed a previously reported approach (DO-MS) for evaluating and optimizing MS experiments.^12^ Specifically, Figures 2 and 4 were generated by plotting variables reported by MaxQuant. The Pearson correlation values displayed in Figure 5(a) were computed from the subset of peptides observed in all samples. For each sample, the correlation was computed between the vector of single-cell Jurkat / U937 reporter ion ratios (mean taken over the 3 replicates of each cell-type before computing each ratio) to the corresponding vector of ratios estimated from the isobaric carrier ratios (Jurkat / U937 reporter ion ratios). Error bars were calculated by repeated sampling with replacement a subset of the peptides. The analysis for Figure 5(b) was performed the same way, but sub-setting first for those peptides with single-cell reporter ion intensity > 2000. To control for the shape of RI distributions in each sample, we sampled subsets as follows: The range of reporter ion values > 2000 was divided into 10 bins of width 0.24 on a log_10_-scale starting at log_10_(2000), and then one peptide was sampled from each bin. The correlations were computed using the sampled subsets of peptides with reporter ion intensities having similar distributions.

## Supplemental website

scope2.slavovlab.net/mass-spec/Isobaric-carrier-optimization

## Acknowledgments

We thank R. Gray Huffman, Edward Emmott, and Prof. Barry Karger for constructive comments. This work was funded by a New Innovator Award from the NIGMS from the National Institutes of Health to N.S. under Award Number DP2GM123497, an Allen Distinguished Investigator award through The Paul G. Allen Frontiers Group to N.S., a CZI Seed-networks grant from the Chan Zuckerberg Initiative to N.S., and through a Merck Exploratory Science Center Fellowship, Merck Sharpe & Dohme Corp. to N.S.

## Competing Interests

The authors declare that they have no competing financial interests.

## Supplemental Figures

**Figure S1.**
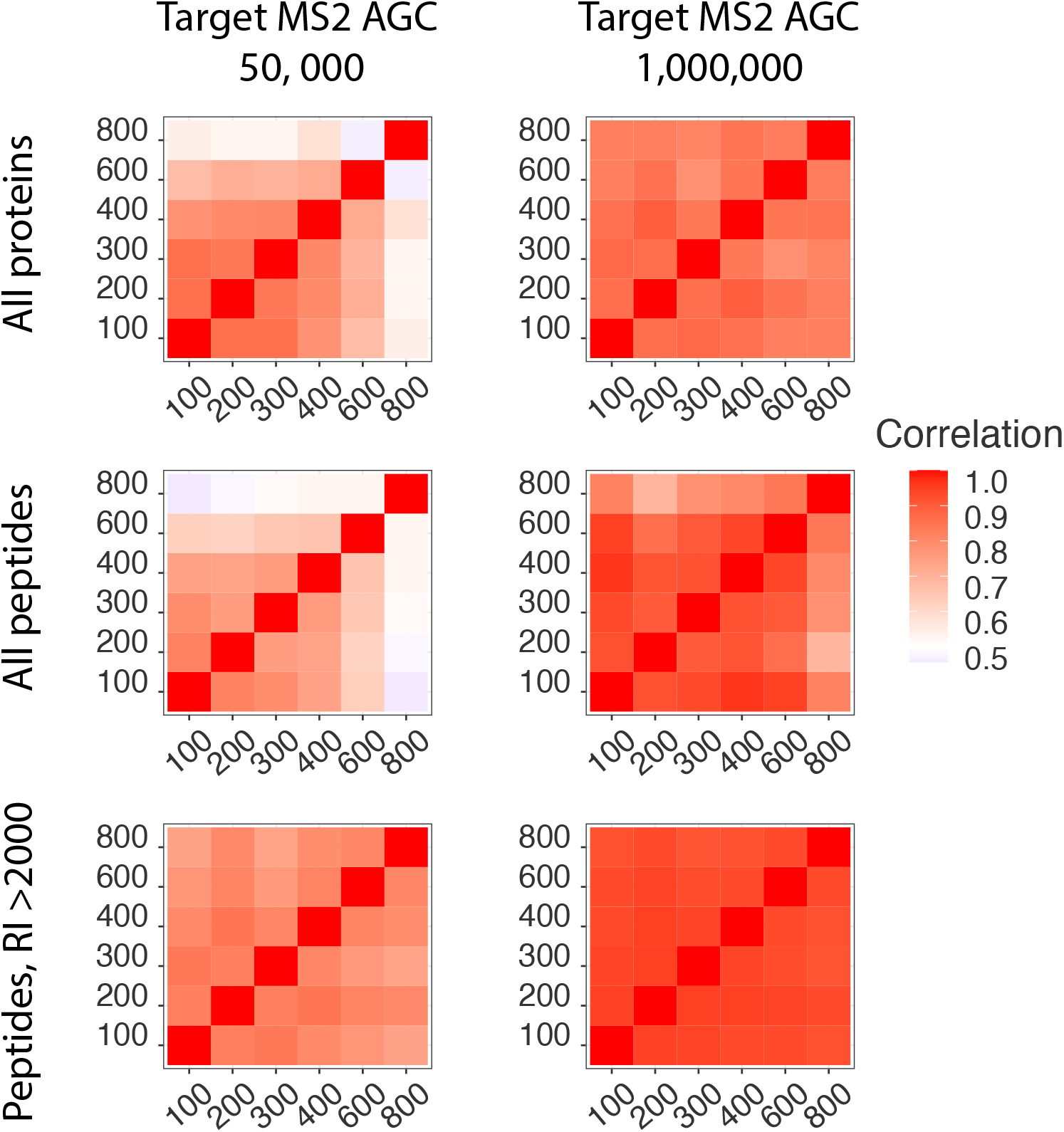
Correlations between protein fold-changes estimated from the single-cell channels of standards with different number of cells in the isobaric carrier. Correlations (Pearson) are computed using peptides in common to every experiment and peptides in common to every experiment with a mean reporter ion intensity of > 2000 in the single cells across two levels of Target MS2 AGC. In this analysis, correlations are computed between single cell quantitative values, as opposed to between single cell quantitative values and bulk values, as in Figure 7.

## Notes

http://scope2.slavovlab.net/

### Competing Interest Statement

The authors have declared no competing interest.

### Summary of Updates

We added an additional figures and analysis reporting on the number of peptide fragments per peptide. We also added a new figure 8 showing the effects of directly decreasing or increasing ion accumulation times on missing data and quantification accuracy. There is no change in the principal findings and guidelines.

